# Amyloid β interaction with membranes: Removal of cholesterol from the membranes to catalyze aggregation and amyloid pathology

**DOI:** 10.1101/2025.08.20.671368

**Authors:** Rishiram Baral, Ruan van Deventer, Yuri L. Lyubchenko

**Author notes:** Corresponding author: Yuri L. Lyubchenko.

## Abstract

The interplay between the cholesterol metabolism and assembly of Aβ42 (the 42- residue form of the amyloid-β peptide) peptides in pathological aggregates is considered as one of the major molecular mechanisms in development of Alzheimer’s disease (AD). Numerous in vitro studies led to the finding that the high cholesterol levels in membranes accelerate the production of Aβ aggregates. The molecular mechanisms explaining how cholesterol localized inside the membrane bilayer catalyzes the assembly of Aβ aggregates above the membrane remain unknown. We addressed this problem by combining different AFM modalities, including imaging and force spectroscopy, with fluorescence spectroscopy. Our combined studies revealed that Aβ42 was capable of removing cholesterol from the membrane. Importantly, physiologically low concentrations of Aβ42 demonstrate such ability of monomeric Aβ42. Extracted cholesterol interacts with Aβ42 and accelerates its on-membrane aggregation. We propose a model of interaction of Aβ42 with membranes based on the ability of Aβ42 to extract cholesterol, which explains several AD associated observations related to cholesterol interplay with Aβ42 aggregation resulting in the AD onset and progression.

---

The age-related formation and accumulation of Amyloid beta (Aβ) aggregates in the extracellular space of the brain, forming plaques is a characteristic hallmark of Alzheimer’s disease (AD) ^1,2^. Elucidating the molecular mechanisms by which intrinsic and extrinsic factors influence the aggregation of Aβ42 is critical for devising effective therapeutic interventions that target and inhibit its self-assembly^3,4^. Physiological concentration of Aβ42 in the low nanomolar level is the factor preventing the assembly of Aβ aggregates in test tubes, yet lipid membranes can affect the aggregation of Aβ ^5–7^. We have shown that membranes are capable of catalyzing of Aβ aggregation at physiologically relevant low nanomolar concentrations of the peptide further supporting a critical role of cellular membranes in the formation of pathological amyloid aggregates ^8,9^. This is the *on-surface aggregation* pathway allowing for a spontaneous aggregation of Aβ peptides of different sizes and α-synuclein protein at the nanomolar concentration range ^8^. The process takes place at ambient conditions, physiological pH values, and without additional mechanical stimulation of aggregation. We developed a theoretical model for the surface-mediated catalysis and tested the model in experiments ^9^. According to the model, aggregation starts with protein monomers transiently attaching to the surface due to molecular interactions. This process increases the local concentration of proteins, which in turn increases the probability of oligomerization reactions to occur on the surface. Of note, the catalytic effect of surfaces in amyloid aggregation explains the experiments on aggregation of Aβ40 in cell culture at low nanomolar concentrations^10^. Our results are in line with ^11^ that reported catalytic properties of the zwitterionic lipid vesicles during the formation of Aβ42 fibrils (reviewed in ^12–15^).

The membrane composition is an important factor contributing to the on-membrane aggregation of Aβ42 which was documented by various methods. Note a study ^16^ in which the role of cholesterol in accelerating Aβ42 aggregation in the presence of vesicles composed of cholesterol and other phospholipids was demonstrated. The lipid metabolism and alteration of lipid composition in the brain are strongly associated with the pathogenesis of AD ^17^.

Cholesterol is a fundamental constituent of cellular membranes and plays a pivotal role in the nervous system ^18^. The brain, as one of the organs with the richest cholesterol content in the human body, accounts for only 2 % of total body weight but contains 25 % of total body cholesterol ^19–21^. Alterations in brain cholesterol metabolism are related to AD and other neurodegenerative disease ^22,23^. The reduction in the efficiency of lipid transport mechanisms with age can lead to lipid imbalances and contribute to neurodegeneration ^24^. Suppression of cholesterol synthesis in astrocytes substantially reduces Aβ42 aggregation. Initial investigations into AD’s lipid-related mechanisms have revealed disrupted cholesterol movement from astrocytes to neurons clearly showing the interrelationship between the cholesterol homeostasis and AD ^24^. Evidence from the literature links AD pathogenesis with an increased concentration of cholesterol ^1,16,25^. It was demonstrated that membrane cholesterol content plays a key role in the neurotoxicity of β-amyloid ^26^.

The membrane composition is the factor defining the catalytic property of the membrane, and the membrane bound-cholesterol significantly accelerates the Aβ42 aggregation at physiologically relevant low nanomolar protein concentration ^27,28^. The Aβ42 aggregation on vesicles containing cholesterol revealed that the kinetics of aggregation depend on the cholesterol concentrations^27,28^ but the molecular mechanism explaining how cholesterol in membranes facilitates aggregation of amyloids above the membrane remains unknown.

Here, we aim to unravel this mechanism by thoroughly characterizing the interaction between Aβ42 and cholesterol-containing membranes. The studies with the use of various AFM modalities and fluorescence experiments revealed that Aβ42 is capable of extracting cholesterol from membranes, which leads to accelerated aggregation. The implication of this phenomenon to understanding the interplay between the cholesterol homeostasis, membrane stiffness and Aβ42 pathology is discussed.

## Results and Discussion

### Cholesterol in the membrane catalyzes Aβ42 protein aggregation

We assembled supported-lipid bilayers composed of 0.25 mg/ml 1-palmitoyl-2-oleoyl-sn-glycero-3-phospho-L-serine (16:0-18:1 POPS) and 1-palmitoyl-2-oleoyl-glycero-3-phosphocholine (16:0-18:1 POPC) in a 1:1 molar ratio with different cholesterol concentrations. Solution of Aβ42 was placed above the bilayer and Atomic Force Microscopy (AFM) imaging and force spectroscopy was applied to characterize aggregation of Aβ42 monomers and its effect on the bilayer. The experiments were conducted using 50 nM Aβ42 peptides, which is in the range of physiologically relevant concentrations.

Three different concentrations of cholesterol (10%, 20%, and 30%) were used to generate stable supported-lipid bilayers with a topographically smooth morphology as described in the methods section. The formation of the bilayer was verified by measuring the thickness of the bilayer at the edge of a smooth area (white dashed line in Figure S1A) and the results from multiple measurements (n=100) (Figure S1B) produce the mean value 4.23 ± 0.92 nm, which corresponds to the typical thickness of similar bilayers^29,30^. The smoothness of the bilayers was monitored with AFM, by measuring the root-mean-square (RMS) roughness of the smooth area. The smoothness of the entire area produced mean values of 0.059 ± 0.0017 nm, which is suitable for detecting the formation of amyloid aggregates^28–30^. This parameter remains constant during the 6-hour observation time (Figure S1C and Table S1)

Once the bilayer was assembled, 50 nM Aβ42 solution in 10 mM HEPES buffer with 150 mM NaCl and 10 mM CaCl_2_ was added on top of the bilayer, and the assembly of aggregates was monitored with time-lapse AFM imaging. The set of data characterizing the Aβ42 aggregation on bilayers without and with 10%, 20%, and 30% of cholesterol taken after 6 hours of continuous observations is shown in Figure 1. Selected AFM images in Figures A, B, C and D demonstrate the increase of number aggregates as the cholesterol concentration increases. A set of AFM images corresponding to different times for all bilayers is displayed in (Figure S2). The aggregates appear as bright globular features in all bilayers, which is consistent with our previous paper ^27,28^, and the number of aggregates (Figure 1E) and their sizes (Figure 1F and Figure S3)) grow with the increasing cholesterol concentration as a function of time.

**Figure 1.**
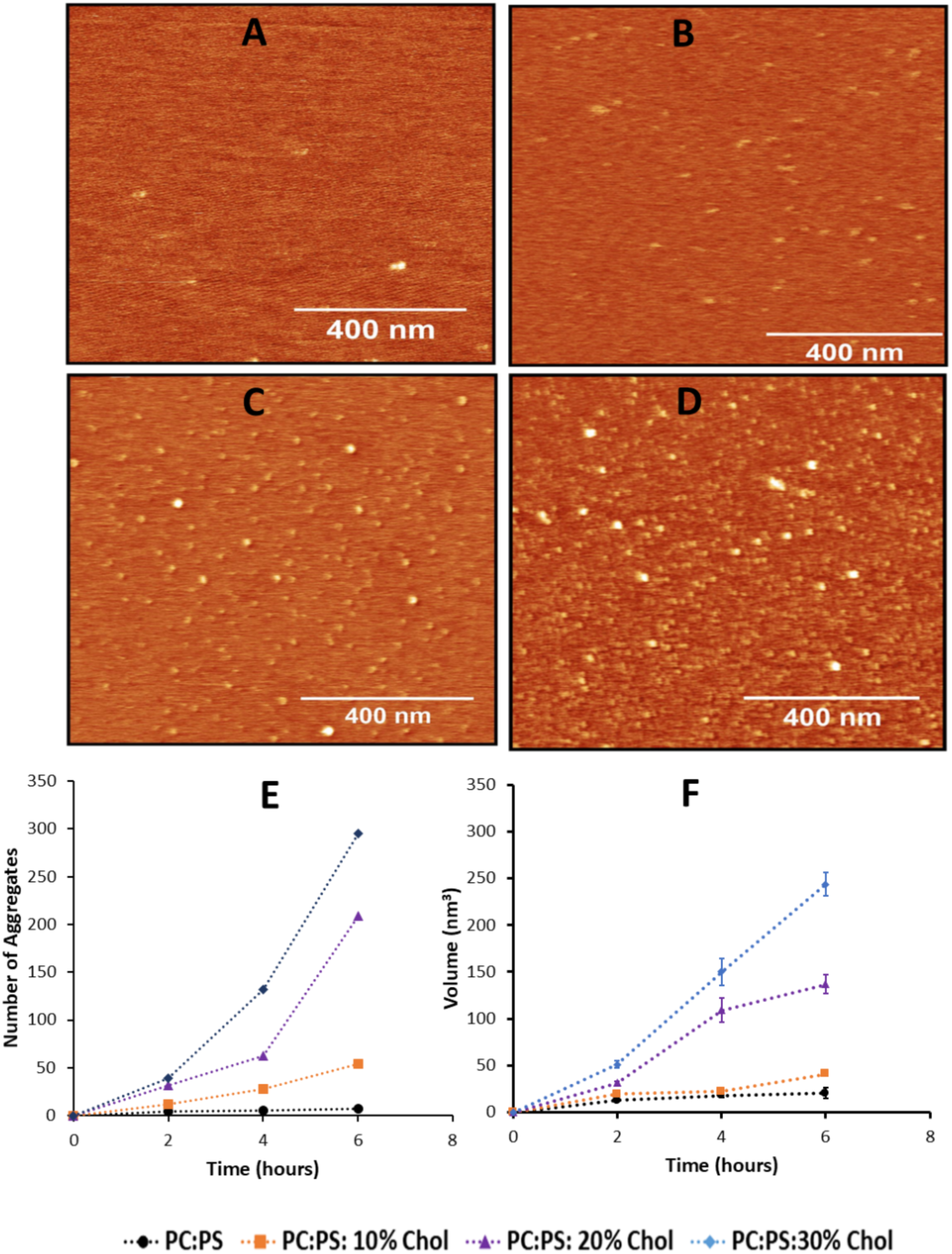
AFM data for aggregation of 50 nM Aβ42 on POPC: POPS bilayer depending on cholesterol in the bilayer. (A-D) AFM images of the aggregation of 50 nM Aβ42 on 0.25mg/mL POPC: (A) POPS lipid bilayer at 6 hours without cholesterol, (B) with 10% cholesterol, (C) with 20% cholesterol, (D) with 30% cholesterol. The scan size is 1 x 1 µm. (E) The average number of aggregates, (F) the average volume of these aggregates over time. The number of aggregates values represents numbers of aggregates in 1 x 1 µm scan of the same size; the volumes are the mean volume of the aggregates with error bars as error of the mean.

### 2.2. Aβ42 reduces membrane stiffness

We assessed how Aβ42 aggregation influences the mechanical properties of bilayers containing 10%, 20%, and 30% cholesterol. During this process, two parameters of the membrane were measured: stiffness and thickness. Stiffness was characterized by Young’s modulus, with multiple measurements performed in parallel with topographic imaging, ensuring that mechanical properties were recorded for each image. Mean Young’s modulus values were calculated from histograms composed of hundreds of measurements per time point. The respective histogram plots for different cholesterol concentrations are shown in Figure S4, and the mean values generated from these histograms are plotted in Figure 2. The results demonstrate that cholesterol increases the initial stiffness of the bilayers, with the highest value observed at 30% cholesterol (Figure 2A, blue points).

**Figure 2.**
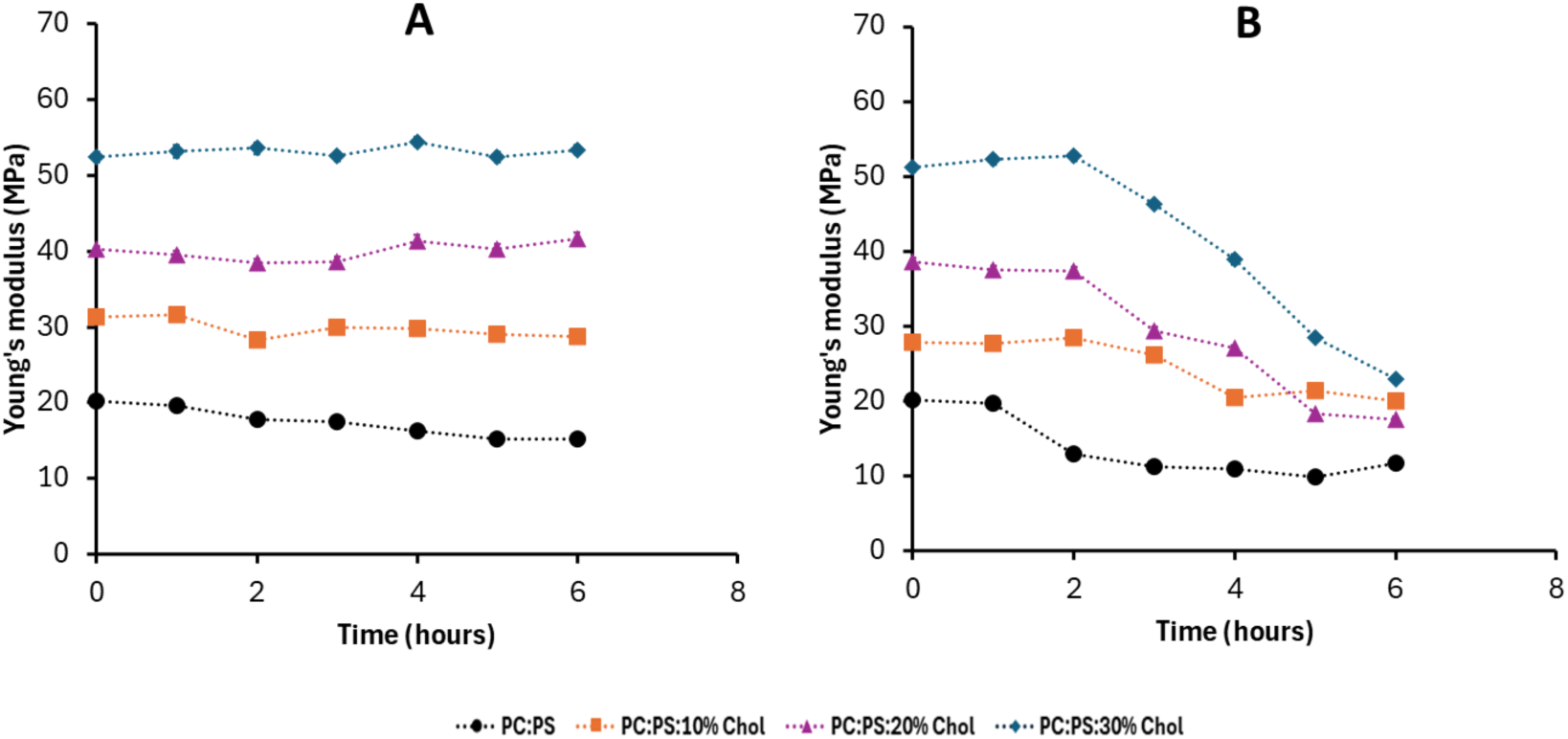
Young’s modulus of POPC: POPS bilayer with different concentrations of cholesterol with and without 50 nM Aβ42 above the bilayer. Young’s modulus of 0.25 mg/mL phospholipid bilayer with different mol% of cholesterol from 0 hour to 6-hour. (A) In the absence of 50 nM Aβ42 (n=100). (B) In the presence of 50 nM Aβ42 from 1 h to 6 h (n=100). Multiple force points taken at various locations within the 1 x 1 µm surface were obtained and 100 such points were approximated with Gaussians. The mean values obtained from the approximations along with the standard error of the mean are shown as scattered plots.

Next, Aβ42 solution was placed on the bilayers with the same concentrations of cholesterol and Young’s modulus were measured as function of time. Histograms for Young’s modulus measurements were taken at different times fitted with Gaussians and the data corresponding to 0, 3 and 6 hours are shown in Figure S4. The maxima values of the histograms for bilayers with different concentrations of cholesterol are shown in Figure 2B. These data demonstrate that upon incubation with Aβ42, Young’s modulus values decrease over time for all bilayers, converging to similar values after 6 hours. The strongest effect is for the bilayer with 30% cholesterol concentration (Figure 2B) in which the initial value 54 MPa value drops after 6 hours to ∼20 MPa corresponding to the bilayer stiffness without cholesterol. Control experiments without Aβ42 showed no change in the membrane stiffness (Figure 2A). These findings suggest that in addition to forming aggregates on the bilayer surface, Aβ42 reduces bilayer stiffness.

Along with the stiffness measurements, the thickness of membranes as a function of time was measured. We used breakthrough force methodology in which the thickness of the bilayers is measured by puncturing it with an AFM tip ^31,32^. These measurements were performed in parallel with stiffness measurements for the POPS: POPC bilayers containing 20% cholesterol and the results are shown in Figure 3. A topographic AFM image of a 1 × 1 μm area, where breakthrough- force measurements were taken at different time points is shown in Figure 3A (at 0 hour), where the image depicts the surface at 0 hour. Force spectroscopy curves collected from various locations across the AFM-defined surface are overlaid and representative image is shown in Figure 3B, depicting the breakthrough force curve at 0 hour. Multiple force curves were taken at various locations within the surface, and 100 such curves were overlaid in each frame. The horizontal distance between the minimum and maximum on the force curve produces the thickness of the bilayer (∼5 nm), and the maximum of the force in the profile corresponds to the force value required for penetrating the bilayer (∼2.5 nN). The overlay of force curves illustrates the uniformity of these parameters over the entire image. The corresponding thickness values were recorded at each hour between 1 and 6 hours and are compiled as histograms are shown in Figure S5. The histograms were fitted with Gaussians and the mean values from these histograms, plotted in Figure 3C. The scattered points remain constant within a range of ∼4 nm, indicating no significant change in bilayer thickness in the presence of Aβ42.

**Figure 3.**
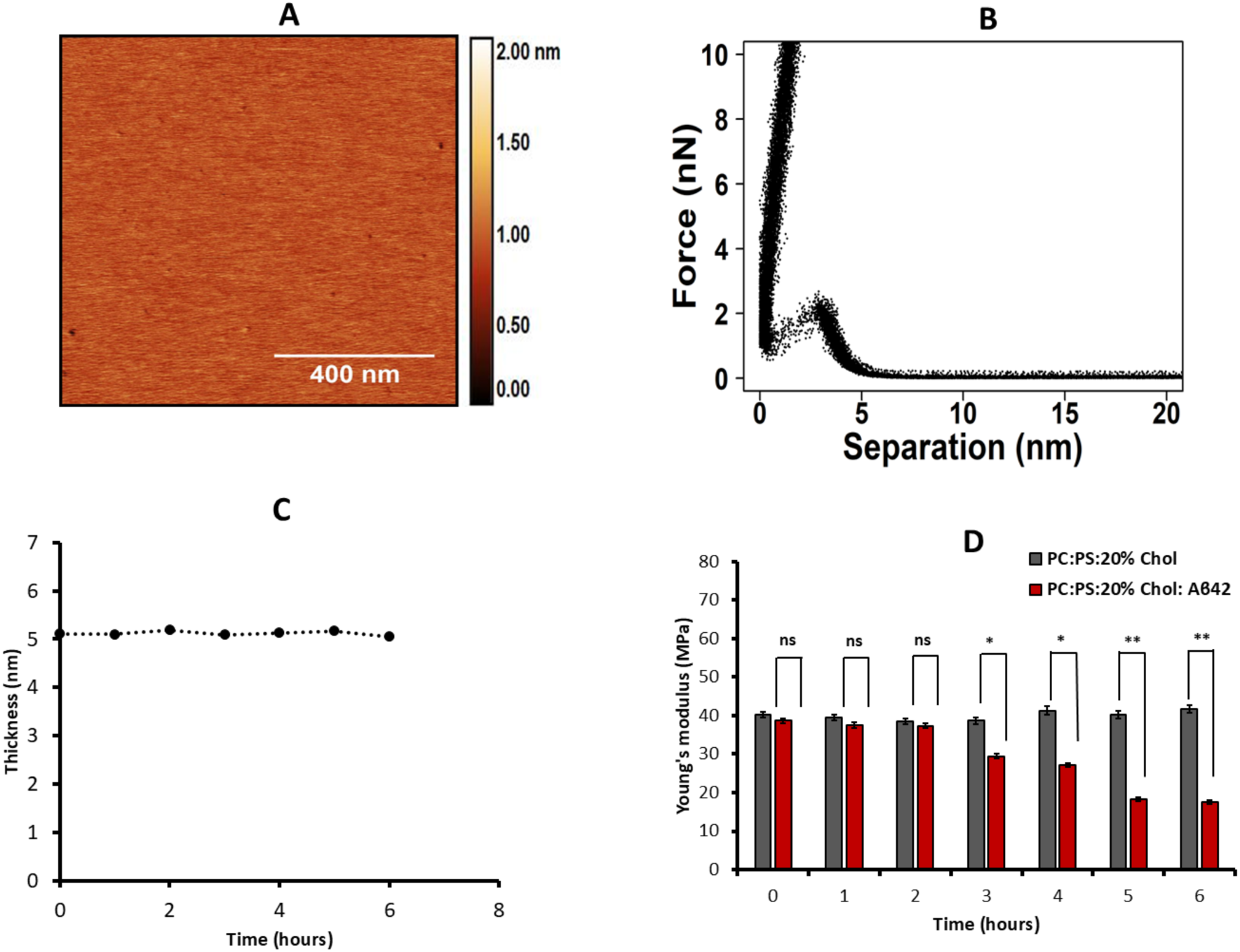
Breakthrough force spectroscopy and thickness for POPC:POPS with 20% Chol bilayer incubated with Aβ42 on top of the bilayer. (A) AFM image of 0.25 mg/mL POPC:POPS:20% Chol phospholipid bilayer incubated with Aβ42 where the breakthrough force was measured. The scan size is 1 x 1 µm. (B) The force-separation profiles correspond to the images shown in Figure 3A. (C) Average thickness of the membrane obtained from the force-distance curve from the 0-time point to 6 hours. Multiple thickness values were measured at several areas within the 1 x 1 µm surface, and 100 such points were approximated with Gaussians. The mean values obtained from the approximations, along with the standard error of mean, are shown as bar graph. (D) Young’s modulus of 0.25 mg/mL phospholipid bilayer with 20% of cholesterol measured on the surface shown in Fig. 3A from 0 hour to 6 hours. Multiple Young’s modulus values taken at various locations within the 1 x 1 µm surface were obtained and 100 such points were approximated with Gaussians. The mean values obtained from the approximations, along with the standard error of mean, are shown as a bar graph.. The difference in means between both groups were tested through student t-test, with significance levels as *p < 0.05, and **p < 0.01.

Parallel to these measurements, Young’s modulus measurements were done as described above. The histograms were approximated with Gaussians (as shown in Figure S6) and the results for the mean values are plotted as bars in Figure 3D. Similar to the data in Figure 2C, the red bars values decrease with the incubation time reaching after 6 hours value close to the one for POPC/POPS bilayer without cholesterol. Gray bars correspond to the control experiments with the same bilayer without adding Aβ42, no changes in the stiffness were obtained for the control.

### 2.3. Depleting cholesterol from membranes

We hypothesize that the drop in bilayer stiffness to values similar to those without cholesterol is due to the removal of cholesterol. To test this hypothesis, we conducted force spectroscopy studies using methyl β-cyclodextrin (MβCD), a known cholesterol-extracting agent ^33,34^.

A solution of MβCD (3 mM in 10 mM HEPES buffer (pH 7.4) containing 150 mM NaCl and 50 mM CaCl_2_) was placed on the top of a POPC:POPS bilayer with 20% cholesterol and changes in Young’s modulus were monitored for a time period over 6 hours. The mean Young’s modulus values are shown in (Figure 4A, orange data points), with the error bars indicating the standard deviation. Within the first 15 minutes of MβCD exposure, the bilayer stiffness decreased from 35.3 ± 4.6 MPa to 18.2 ± 3.2 MPa, (shown in zoomed circle of Figure 4A) with a gradual decrease over the entire observation period. The final Young’s modulus value matches the stiffness of the POPC: POPS without cholesterol (Figure 2A, Table S2).

**Figure 4.**
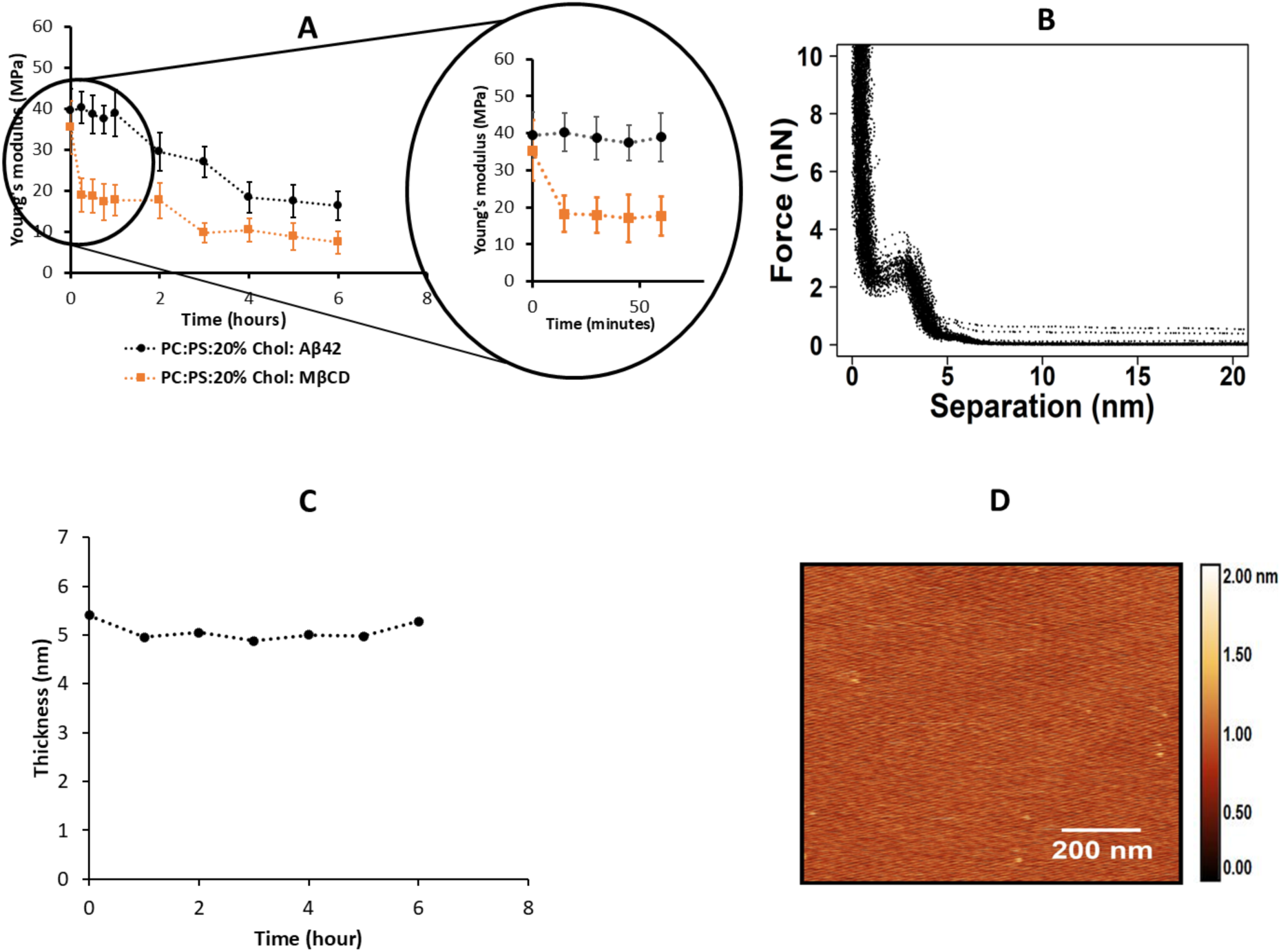
Breakthrough force spectroscopy and thickness for POPC:POPS with 20% Chol bilayer incubated with MβCD on top of the bilayer. (A) Young’s modulus of 0.25 mg/mL phospholipid bilayer with 50 nM Aβ42 (black dots) and 3 mM MβCD (orange dots) (n=100). The zoomed circle on the right illustrates the kinetics for the one hour of observation. (B) The force-distance profiles corresponding to the images shown in Fig. 4D. Multiple force curves were taken at various locations within the surface, and 100 such curves were overlaid in each frame. The horizontal distance between the minimum and maximum on the force curve produces the thickness of the bilayer (∼5 nm), and the maximum of the force in the profile corresponds to the force value required for penetrating the bilayer (∼4 nN). (C) Thickness values of the membrane exposed to 3 mM MβCD depending on time. The mean values obtained from the approximations, along with the standard deviations, are shown as bar graph. (D) AFM image of 0.25 mg/mL POPC: POPS:20% Chol phospholipid bilayer at 6 hours showing the area where the breakthrough force was measured. The scan size is 1 x 1 µm.

The bilayer thickness was measured in conjunction with Young’s modulus. A subset of the force curves is shown in Figure 4B, and corresponding histogram of thickness for each time point are assembled in Fig S7. The mean thickness values over time, plotted in Figure 4C, indicate that the bilayer thickness remains unchanged despite the reduction in stiffness. An AFM image (Figure 4D) taken after 6 hours of MβCD exposure shows that the bilayer surface remains topographically smooth.

For direct comparison, we repeated the experiment for a similar bilayer for 50 nM Aβ42. The data in (Figure 4A, in black symbols) show the gradual decrease in Young’s modulus. Comparing two sets of data (Figure 4A) reveals that MβCD and Aβ42 exhibit a similar effect on the mechanical properties of the bilayer. Importantly, MβCD and Aβ42 reduce the bilayer stiffness to levels comparable to cholesterol-free membranes. This supports our hypothesis that Aβ42 may deplete cholesterol from the bilayer through a mechanism analogous to that of MβCD.

### 2.4. Extraction of cholesterol from the membrane by Aβ42 and MβCD: fluorescence spectroscopy studies

To directly test the hypothesis on the cholesterol extraction by Aβ42 and detect free cholesterol depleted from the membrane, we conducted experiments using fluorescently labeled cholesterol incorporated into a POPC: POPS bilayer. If cholesterol is released, it can be detected via fluorescence measurements of the solution above the membrane. We used 25-[N-[(7- nitro-2-1,3-benzoxadiazol-4-yl)methyl]amino-27-norcholesterol (25-NBD cholesterol), a common cholesterol analog, in fluorescence studies ^35^. A POPC: POPS bilayer containing 20% 25-NBD cholesterol was assembled, and 50 nM Aβ42 was placed on the top of the membrane. Aliquots of the solution above the membrane was collected and fluorescence spectra were measured. The set of spectra measured over time shown in Figure 5A demonstrate the increase of fluorescence in the solution and the effect is higher for longer incubation time. Control experiments with the buffer above the bilayer without Aβ42 in Figure 5B shows some increase of fluorescence that can be attributed to the spontaneous release of weakly incorporated cholesterol from the bilayer. Thus, there is a dramatic difference in the fluorescence values for experiments with Aβ42 compared with the control. An apparent increase in intensity of fluorescence spectra over time suggests cholesterol accumulation above the membrane.

**Figure 5.**
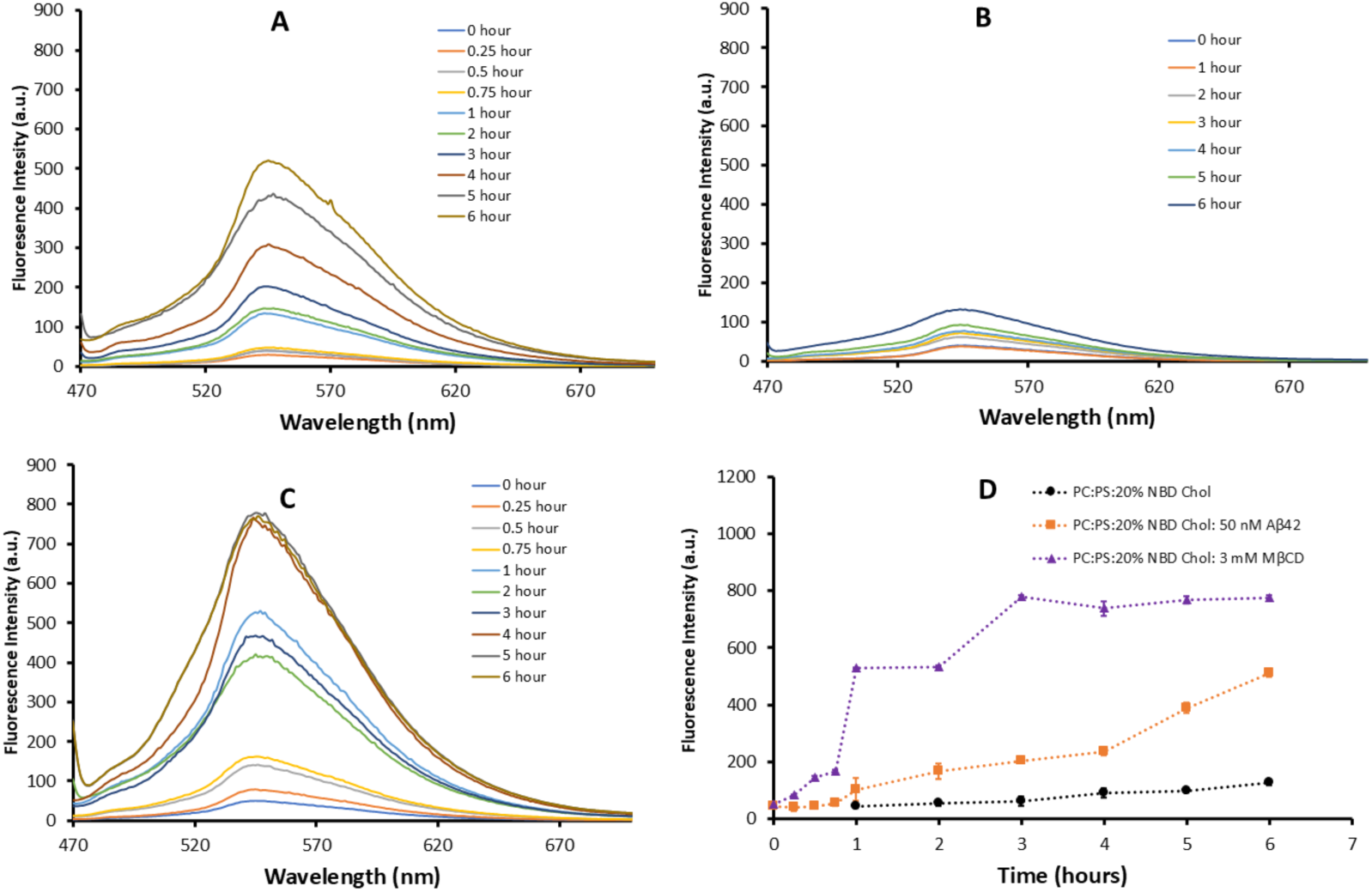
Fluorescence intensity measurements of solution above POPC: POPS bilayers containing 20 mol% fluorescently labeled cholesterol (25-NBD cholesterol). The temporal changes in fluorescence intensity were monitored in solutions collected from POPC: POPS bilayers (0.25 mg/mL) incorporated with 20% 25-NBD cholesterol under the following conditions: (A) incubation with 50 nM Aβ42, (B) exposure to 10 mM HEPES buffer with salt, (C) treatment with 3 mM MβCD (D) mean of the maxima values of fluorescence intensity recorded at each time point for each experimental condition with standard error of the mean were plotted against time. Excitation of the fluorophore was achieved at 450 nm, and emission spectra were recorded from 470 nm to 700 nm, with a peak emission observed at 544 nm.

Similar experiments were performed with MβCD and the results of the fluorescence measurements are shown in Figure 5C. There is a time-dependent increase of fluorescence due to the accumulation of 25-NBD cholesterol in the bulk solution above the membrane. Fluorescence maxima from the fluorescence spectra obtained from three independent experiments (shown in Tables S3, S4 and S5) were plotted as a function of time (Figure 5D). The accumulation of cholesterol occurs in the presence of Aβ42 and MβCD although the rate is faster for MβCD. However, this difference should be considered in the context of the nearly five orders of magnitude lower Aβ42 concentration compared with that of MβCD used in these experiments.

### 2.5 AFM topographic and force spectroscopy studies of Aβ42 interaction with POPC:POPS bilayer containing fluorescently labeled 25-NBD cholesterol

Although 25-NBD cholesterol is commonly used as a cholesterol analog in fluorescence measurements, potential differences between fluorophore-labeled and native cholesterol cannot be excluded. To assess whether the fluorophore moiety affects Aβ42 aggregation on the membrane, we assembled a 0.25 mg/mL POPC: POPS bilayer containing 20 % 25-NBD cholesterol and monitored the aggregation of 50 nM Aβ42 on its surface using AFM for up to six hours. Images taken at six hours (Figure 6B) confirmed aggregate formation on the bilayer compared to the control (Figure 6A).

**Figure 6:**
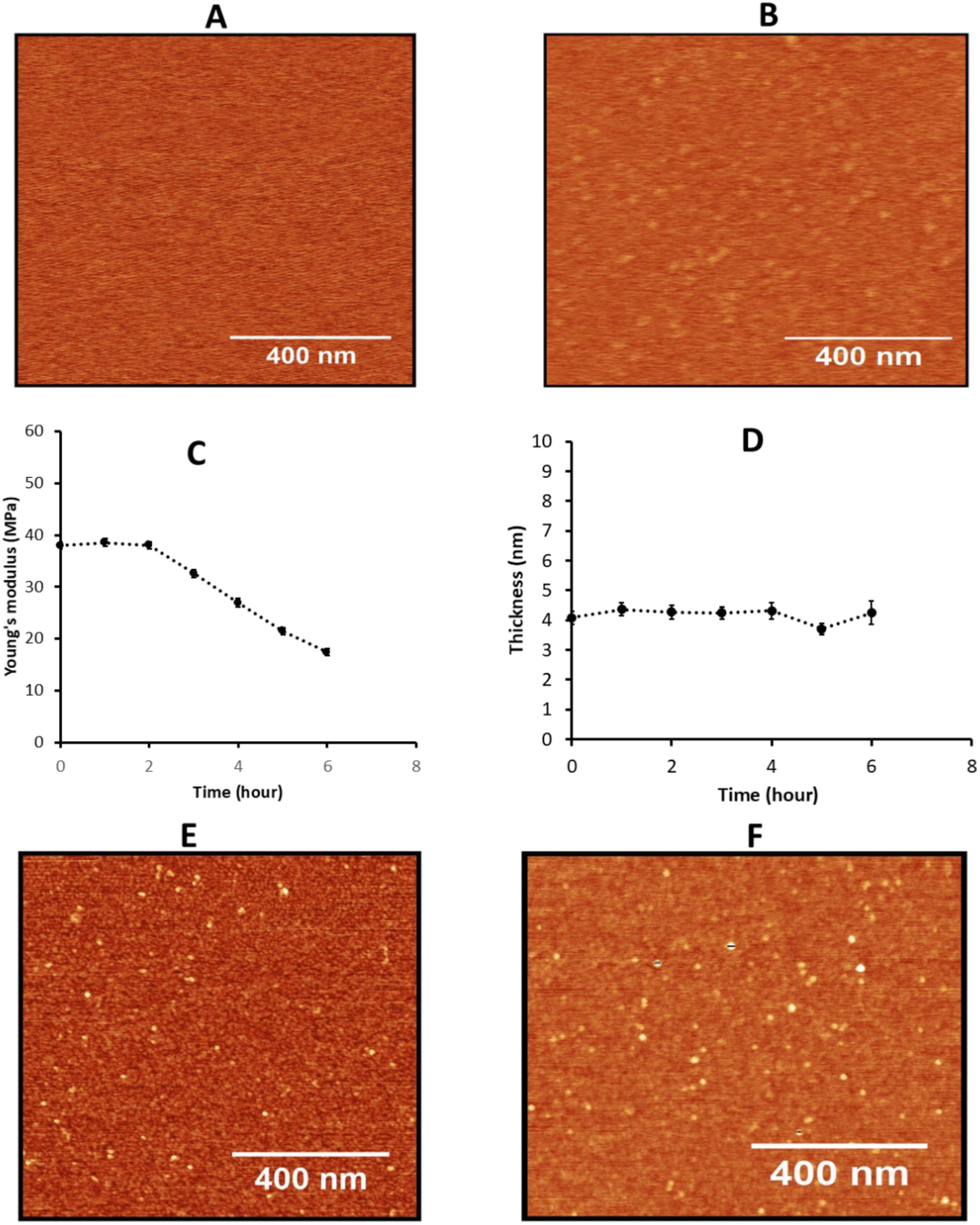
AFM-time lapse imaging and mechanical characterization of bilayer with 25-NBD cholesterol. AFM imaging was performed on a 0.25 mg/mL POPC:POPS phospholipid bilayer incorporating 20% 25-NBD cholesterol in the presence of 50 nM Aβ42, with scans acquired at (A) 0 hour and (B) 6 hours. The scan area was set to 1 × 1 µm. (C) Young’s modulus of the phospholipid bilayer under these conditions was determined (n = 100). (D) membrane thickness variations over time (0 to 6 hours) were quantified using force-distance curve analysis. Multiple measurements were conducted across different regions of the 1 × 1 µm scan area, and 100 data points were fitted to Gaussian distributions. The mean thickness values, derived from these approximations, are presented alongside their respective standard deviations as scattered data points. (E, F) AFM time-lapse imaging was conducted on a dry sample obtained by withdrawing the supernatant from a 0.25 mg/mL POPC:POPS phospholipid bilayer incorporating 20% 25-NBD cholesterol. Images corresponding to time points (E)1 hour and (F) 6 hours post-solution removal were analyzed.

Young’s modulus measurements (Figure 6C) illustrate the decrease of stiffness from 37.5 ± 7.3 MPa to 16.2 ± 5.6 MPa, closely aligning with values obtained for the POPC: POPS bilayer containing cholesterol (Figure 2A). Bilayer thickness (Figure 6D) remained unchanged upon Aβ42 incubation, with values comparable to those of cholesterol-containing bilayers. Aliquots from the bulk solution above the membrane were collected, deposited on APS mica, dried, and imaged, and the data were analyzed to characterize additional Aβ42 aggregation (Figure 6E and 6F). Similar to previous findings ^27,28^Aβ42 aggregates accumulate in the solution above the POPC: POPS:25- NBD cholesterol membrane. These results confirm that both cholesterol types; regular (Figure S8) and fluorescently labelled 25-NBD cholesterol (Figure 6) have a similar effect on Aβ42 aggregation on membranes.

### 2.6. Discussion

The results described above revealed novel features of the interaction of Aβ42 with the cholesterol containing phospholipid bilayers. The topographic AFM studies directly visualized aggregation of Aβ42 on the surface and the process is accelerated by cholesterol (Figure 1), which is in line with our previous finding ^36^. Parallel experiments with the AFM force spectroscopy reveled that the bilayer stiffness in the presence of Aβ42 drops and the effect increases with the incubation time (Figure 2). Importantly, the stiffness of the bilayers over time approaches the values corresponding to the membrane stiffness without cholesterol. Similar results with the bilayer stiffness loss were obtained with the use of the well-known cholesterol extractor MβCD, suggesting that Aβ42 actively extracts cholesterol from the bilayer. This novel key finding on the ability of Aβ42 actively extracts cholesterol is directly demonstrated using a fluorescently labeled cholesterol analog (25-NBD cholesterol). The fluorescence spectra of the solution above the bilayer reveal the accumulation of NBD cholesterol after its release from the bilayer (Figure 5). The cholesterol extraction is the time-dependent process and the effects of cholesterol extraction is close to the one measured for MβCD. Although MβCD extracts cholesterol more rapidly, it was used at a concentration five orders of magnitude higher than Aβ42. After 6 hours, Aβ42 achieved ∼75% of the total MβCD-mediated cholesterol release, highlighting its remarkable efficiency.

Computational modeling ^37^ suggests that MβCD facilitates cholesterol extraction through partial insertion into the membrane, allowing one of its rings to interact with the hydroxy group of the embedded cholesterol with an interaction distance of ∼0.2 nm. In line with this model, recent MD simulations and *in vitro* fluorescence quenching experiments ^38^, have shown that Aβ42 can partially insert into membranes, suggesting that similar to MβCD, a partially inserted Aβ42 may extract cholesterol. Neutron scattering experiments on POPC bilayer with cholesterol revealed the displacement of cholesterol towards the membrane surface in the presence of Aβ42^39^ providing additional evidence to the extraction capability of Aβ42.

Based on these findings, we propose the model for the Aβ42 interaction with membrane shown in Figure 7.

**Figure 7:**
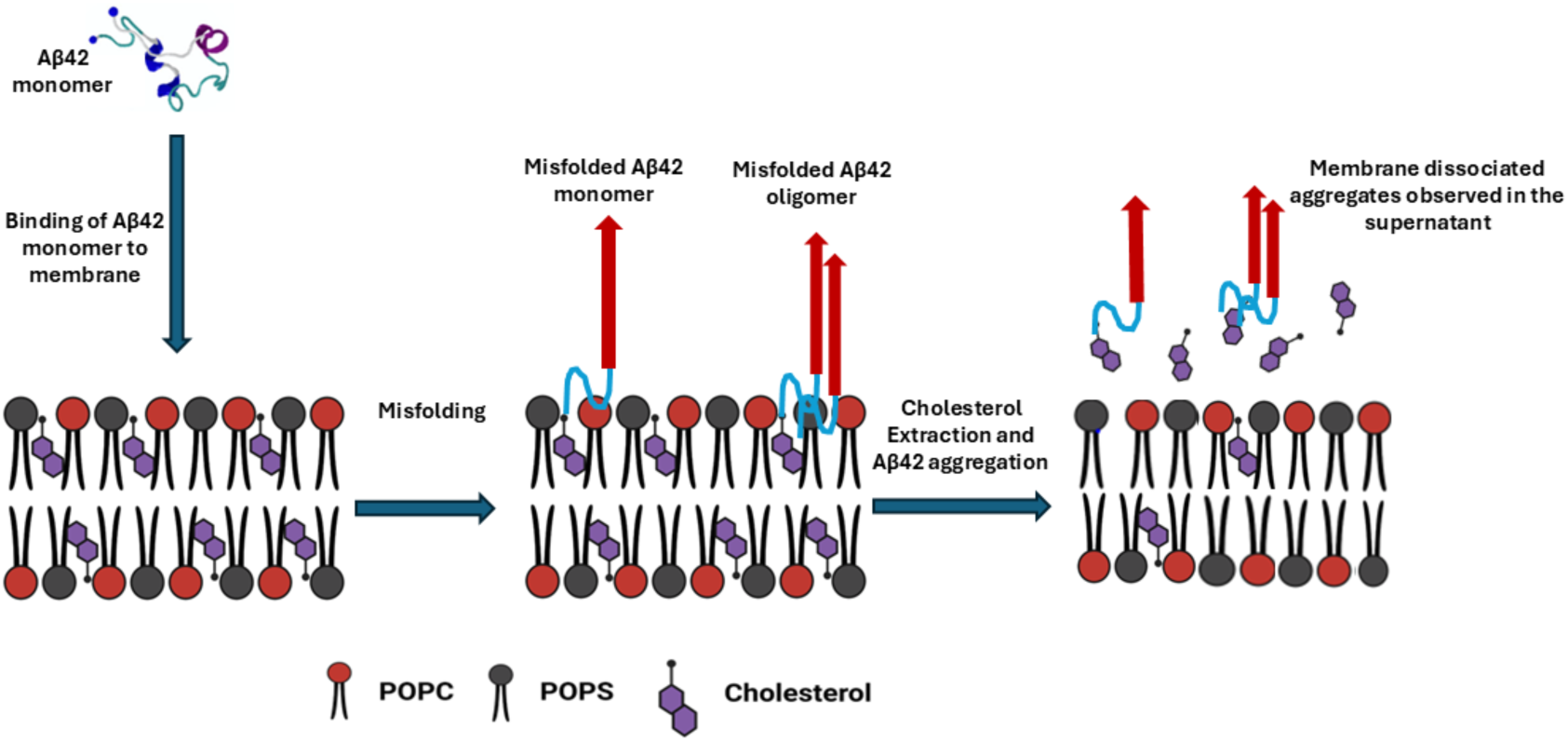
Model for interaction of Aβ42 with membrane: Depletion of cholesterol and the disease associated effects. Membrane binding induces conformational misfolding of monomeric Aβ42, promoting its transition to a pathogenic state. The misfolded protein facilitates cholesterol extraction from the membrane while concurrently assembling into amyloid aggregates. Both the disruption of cholesterol homeostasis and the accumulation of Aβ aggregates synergistically exacerbate Alzheimer’s disease pathology. [Figure created with BioRender.com]

According to this model, Aβ42 after adsorption to the membrane surface and possible misfolding (step 1), extracts cholesterol from membrane (step 2). Extracted cholesterol, accelerates aggregation of Aβ42 above the bilayer (step 3), which is in line with our recent publication ^27^, demonstrating that Aβ42 aggregation kinetics are significantly enhanced in the presence of free cholesterol. Computational modeling presents in the same study ^27^ reveals that the interaction of cholesterol with Aβ42 triggers a conformational switch, promoting its aggregation. This model explains the presence of cholesterol within Aβ42 aggregates and, ultimately, in extracellular plaques, as reported previously ^40,41^. The model and our findings provide explanations to a number of events associated with the development of Alzheimer’s disease.

To contextualize our findings, it is important to consider the physiological and pathological roles of cholesterol in the brain. Although the brain is rich in cholesterol, it does not appear in the free form. The major pool of cholesterol (70%-80%) is found in the myelin sheaths, with the rest of the lipids being in plasma membranes ^42^. Cholesterol is a highly functional molecule in the brain, so its concentrations are tightly regulated ^18^. To this point, the dysregulation of cholesterol homeostasis is often associated with AD development, with its accumulation being one of the risk factors associated with disease development and progression ^43^. For instance, *in vivo* studies (e.g.,^44^) have shown that the intracellular accumulation of cholesterol in microglia induces maladaptive immune responses (such as cholesterol crystal formation, phagolysosomal membrane rupture, and stimulated inflammasomes), which impede remyelination and tissue regeneration. Cholesterol extraction by Aβ peptide contributes to this pathological effect of cholesterol.

Additionally, both *in vitro* and *in vivo* studies have shown that cholesterol accumulates within the mature senile amyloid plaques of AD ^40,41^. Here, our findings on cholesterol extraction by Aβ42 offer a mechanistic explanation for the presence of cholesterol within Aβ42 aggregates and, ultimately, in amyloid plaques observed in AD brains. These results support the view that cholesterol dysregulation contributes to AD pathology and may underlie previously observed histopathological features.

Depletion of cholesterol from the membrane leading to softening of membranes is another potentially pathological feature of Aβ42 associated with its interaction with membranes. Previous *in vitro* studies^45^ have shown that depletion of membrane-bound cholesterol results in the degeneration of dendritic spines and, ultimately, synapse loss-a common pathological feature observed in AD ^46^. Cholesterol itself is largely a hydrophobic molecules primarily localized within the cell membranes, where it regulates membrane fluidity and can interact with neighboring lipids and proteins to regulate membrane trafficking or signal transduction ^47^. Cholesterol extraction decreases the membrane stiffness, which can result in deteriorating the function of membrane-bound proteins. Specifically, lower cholesterol levels can inhibit APP (Amyloid Precursor Protein) processing by secretases ^18^. Importantly, cholesterol molecules are not uniformly distributed within the cell membrane but with other lipids form cholesterol rich rafts. Their formation is critical for various membrane functions, including signal transduction ^48^. Aβ42-mediated cholesterol depletion from these membrane domains can impair membrane dynamics and compromise the function of membrane proteins, many of which rely on the structural and mechanical properties of the lipid bilayer. Interestingly, β- and γ-secretases are primarily localized in cholesterol-enriched lipid rafts, so cholesterol depletion can alter their activity. Since these enzymes are responsible for APP cleavage and the production of Aβ peptides, including Aβ42, such changes can either increase or decrease Aβ levels ^49^ both of which can contribute to AD pathology.

The association of cholesterol with Aβ42 aggregates is the factor that needs to be taken into account in the development of efficient anti-amyloid immunotherapy ^50,51^. The presence of cholesterol in natural amyloid aggregates can be a factor contributing to the lack of their binding to antibodies raised to aggregates assembled without cholesterol.

In conclusion, the current view on the molecular mechanism of AD development is associated with the assembly of Aβ peptides in pathologic aggregates, Aβ oligomers specifically. We discovered a novel function of Aβ42 monomers, namely its capability to extract cholesterol from membranes. Interaction of Aβ42 with free cholesterol explains the molecular mechanism for the cholesterol dependent aggregation of Aβ42 on membranes. At the same time, the accumulation of free cholesterol and the increase of membrane fluidity after cholesterol depletion are the factors associated with AD-related pathologies. Thus, additional Aβ42 associated pathologic factors were discovered, and this discovery changes the Aβ pathology paradigm. These findings suggest that therapeutic efforts for AD treatment should be focused on the development of efficient means of preventing or minimizing Aβ interaction with membranes. Moreover, given that the on-membrane catalysis is the mechanism by which Aβ aggregation at physiologically relevant concentrations occurs^52^, the development of approaches affecting this pathology related interactions of Aβ should be the primary ones. Such approaches have the potential to pave the way for innovative combination therapies that could alleviate the suffering of millions of AD patients.

## MATERIALS AND METHODS

### Methods

#### Materials

1-palmitoyl-2-oleoyl-sn-glycero-3-phosphocholine, CID: 850457 (POPC), 1-palmitoyl-2- oleoyl-sn-glycero-3-phospho-L-serine, CID: 840034 (POPS) and 25-[N-[(7-nitro-2-1,3- benzoxadiazol-4-yl)methyl]amino-27-norcholesterol, CID: 810250 (25-NBD Cholesterol) were purchased from Avanti Polar Lipids (Corda International Plc. Alabama, USA). Lyophilized cholesterol (CAS Number: 57885) and Methyl-beta cyclodextrine (MβCD) (CAS Number: 128446-36-6) was obtained from Sigma-Aldrich (Sigma-Aldrich, Inc, St. Louis, USA). Lyophilized Aβ42 was procured from Genscript (Genscript, New Jersey, USA). Chloroform was procured from Sigma-Aldrich Inc. The buffer solution used in this study is 10 mM HEPES, pH 7.4 with 150 mM NaCl and 10 mM CaCl_2_. In liquid AFM probes, MSNL-10 were purchased from Bruker (Bruker, California, USA).

#### Preparation of Aβ42 Protein Solution

Stock solution of Aβ42 was prepared as described in our previous publication^8,28^. Briefly, lyophilized Aβ42 powder (Genscript) was dissolved and sonicated for 5 minute in 100 µL 1,1,1,3,3,3 Hexafluoroisopropanol (HFIP)maintaining the final concentration of 50 µM to break down pre-formed oligomers. This thin film formed after the evaporation of HFIP overnight was reconstituted in DMSO, preserving the original concentration. This stock solution was stored at −20°C until use. An aliquot of the stock solution was dialyzed against 10 mM HEPES buffer (pH 7.4) and the final concentration was measured using NanoDrop at 280 nm. The dialyzed solution was then diluted to 50 nM using 10 mM HEPES buffer (pH 7.4) containing 150 mM NaCl and 10 mM CaCl_2_.

#### Assembly of 0.25mg/mL POPC:POPS supported lipid bilayers with different concentration of cholesterol

Stock solution of 10 mg/mL POPC, POPS and cholesterol were stored at −20 °C. The experimental working solution was prepared as described in the references^53,54^ with slight modification. Stock solutions of POPC, POPS, and cholesterol were mixed, and chloroform was evaporated under nitrogen to form a thin film. The film was resuspended in 10 mM HEPES (pH 7.4) containing 150 mM NaCl and 10 mM CaCl₂, with a final lipid concentration of 0.25 mg/mL. Cholesterol concentrations were adjusted to 10%, 20%, and 30% in separate solutions. These working solutions were stored at 4°C and used at room temperature unless specified.

Supported lipid bilayers (SLB) were prepared following the methodology outlined in our previous publications^28,29^. Briefly, a piece of mica, used as a solid support, was glued to hydrophobic-coated glass slides, and its surface was freshly cleaved before phospholipid incubation. The phospholipid solution with cholesterol was sonicated for 45 minutes and incubated on the mica surface in a saturation chamber at 60°C for 50 minutes. Solvent evaporation during heating was compensated by adding the deionized water as needed. After incubation, the solution was replaced with 10 mM HEPES buffer (pH 7.4) containing 10 mM CaCl₂ and 150 mM NaCl to remove unruptured vesicles. The sample was cooled to room temperature for 10 minutes before imaging.

#### Treatment of assembled SLB with Methyl-β-cyclodextrin (MβCD) to deplete the cholesterol content

The depletion of cholesterol in the assembled SLB was performed as per the method described in the reference^34^ with slight modification. Briefly, MβCD was dissolved in 10 mM HEPES buffer (pH 7.4) to prepare a 100 mM stock solution, which was further diluted to 3 mM using buffer with 10 mM CaCl₂ and 150 mM NaCl for bilayer stability. A POPC:POPS:20% cholesterol bilayer (0.25 mg/mL) was incubated with 150 µL of 3 mM MβCD for 30 minutes at room temperature. After incubation, MβCD was rinsed off, and the bilayer was treated with 50 nM Aβ42 in buffer with salt. Time-lapse AFM imaging was performed to observe Aβ42 aggregation. Force spectroscopy was used to measure the bilayer stiffness and thickness. A control experiment was conducted without Aβ42.

#### Time-Lapse AFM Imaging

Liquid imaging in AFM using buffer solution was carried out in tapping mode using MFP- 3D instrument (Asylum Research, Santa Barbara, CA, USA). The cantilever “E” of MSNL probes (Bruker, Santa Barbara, CA, USA) having nominal spring constant of 0.1 N/m and typical resonance frequency of 7-9 kHz was used for scanning the bilayer surface. The homogeneity and smoothness of the lipid bilayer surface, devoid of any unruptured vesicles was ensured at the beginning of each time lapse-experiment by scanning the larger area of 10 x 10 µm. Then, a smooth bilayer surface of 1 x 1 µm area was selected, followed by the incubation of Aβ42 solution. AFM imaging on the MFP-3D until 6 hour was automated using the integrated MacroBuilder^TM^ software. The software automatically allowed the parking of the cantilever in between each scan. With this, the number of scans was restricted to prevent damage to the bilayer, as repeated probing by the AFM tip is known to cause degradation.

#### Measurement of Young’s Modulus using AFM Force Spectroscopy

The Young’s modulus of the bilayers was calculated following the method described previously^30,55^. The quantitative determination of the stiffness of the bilayer was performed in contact mode using MFP-3D instrument (Asylum Research, Santa Barbara, CA, USA). MSNL E- tip was used to softly probe the bilayer surface by applying the constant force of 200 pN with approach and retract velocity of 400 nm/sec. The Young’s modulus was plotted from force distance curve using DMT model^36^. Young’s modulus value of each point of probing was derived from individuals pixels and plotted as histogram.

#### Measurement of thickness of the bilayer using the breakthrough force spectroscopy

The rupture event of the bilayer was studied as per the method mentioned in the references ^31,32,56^. 0.25 mg/mL POPC:POPS:20% Chol SLB was assembled on freshly cleaved mica as described in method above. The formation of the bilayer was confirmed by measuring the height of the edge of the bilayer with horizontal cross-section accompanied with the measurement of Young’s modulus, using the constant force of 200 pN. Followingly, the bilayer was probed with high force of 10 nN to visualize the potential rupture events. The force was kept constant throughout the experiment until 6 h and the approach velocity was set to 400 nm/sec. Breakthrough force was quantified from force spectroscopy measurements by identifying the peak force value within the recorded force curve. The bilayer thickness was calculated by subtracting the distance at which the force curve begins to ascend from the distance corresponding to the peak of the force curve. The force-distance data obtained was analyzed using RStudio. The breakthrough and thickness data obtained were plotted as histogram and fitted with single peak Gaussian fit using Magicplot Pro 2.3.92.

#### Measurement of fluorescence intensity of 25-NBD cholesterol released from the assembled bilayer

The release of the fluorescently labelled cholesterol from the 0.25 mg/mL bilayer was measured as described previously^33,57,58^ with few modifications. 0.25 mg/mL POPC:POPS bilayer incorporating 20 mol% of 25-NBD-cholesterol was prepared following the protocol as mentioned previously. After assembly, the bilayer was thoroughly rinsed with 10 mM HEPES buffer (pH 7.4) to eliminate unincorporated fluorescently labeled cholesterol. Subsequently, the bilayers were incubated under three experimental conditions: (1) 50 nM Aβ42, (2) 3 mM MβCD, or (3) control buffer containing 10 mM HEPES, 150 mM NaCl, and 10 mM CaCl₂. Aliquots were collected hourly from the bulk solution above the bilayer, and fluorescence intensity was recorded using a Varian Cary Eclipse Fluorescence Spectrophotometer. Fluorescence measurements were conducted with an excitation wavelength of 450 nm and an emission scan range of 470–700 nm, with a peak emission observed at 544 nm.

#### Data Analysis

The AFM images of the lipid bilayer acquired using the MFP-3D were flattened, visualized and analyzed using Gwyddion v2.66 (Gwyddion, Czech Republic). The formation of the phospholipid bilayer was confirmed by measuring the height of the bilayer from the mica surface upto the edge of bilayer. The number of Aβ42 aggregates and their volumes were determined using Enum Feature tool in FemtoScan software (Advanced Technologies Center, Moscow, Russia). The data generated from the AFM topographic and force spectroscopy experiments were plotted as histograms using Origin Pro software (OriginLab, Northampton, MA, USA). When necessary the data were fitted with single-peak Gaussian distribution. The error bars in graphs represent the standard deviation.

#### Data availability

The data supporting the findings of this study are available within the Article and its Supplementary information files.

## Acknowledgements

We thank Lyubchenko Lab members for useful insight.

## Author contributions

The experiments were designed by Y.L.L., and R.B. carried out the experiments. Data analyses were performed by R.B. Rscript for data analysis was customized by R.V.D. Y.L.L., R.B. and R.V.D. did data interpretation and hypothesizing the model. The work was supervised by Y.L.L. The manuscript was written by R.B., R.V.D. and Y.L.L. Funding acquisition was done by Y.L.L. All authors have read and agreed to the published version of the manuscript.

## Supplementary information

The online version contains supplementary material available at……..

**Correspondence and requests for materials** should be addressed to Yuri L. Lyubchenko at.

---

**Supplementary Fig. 1:**
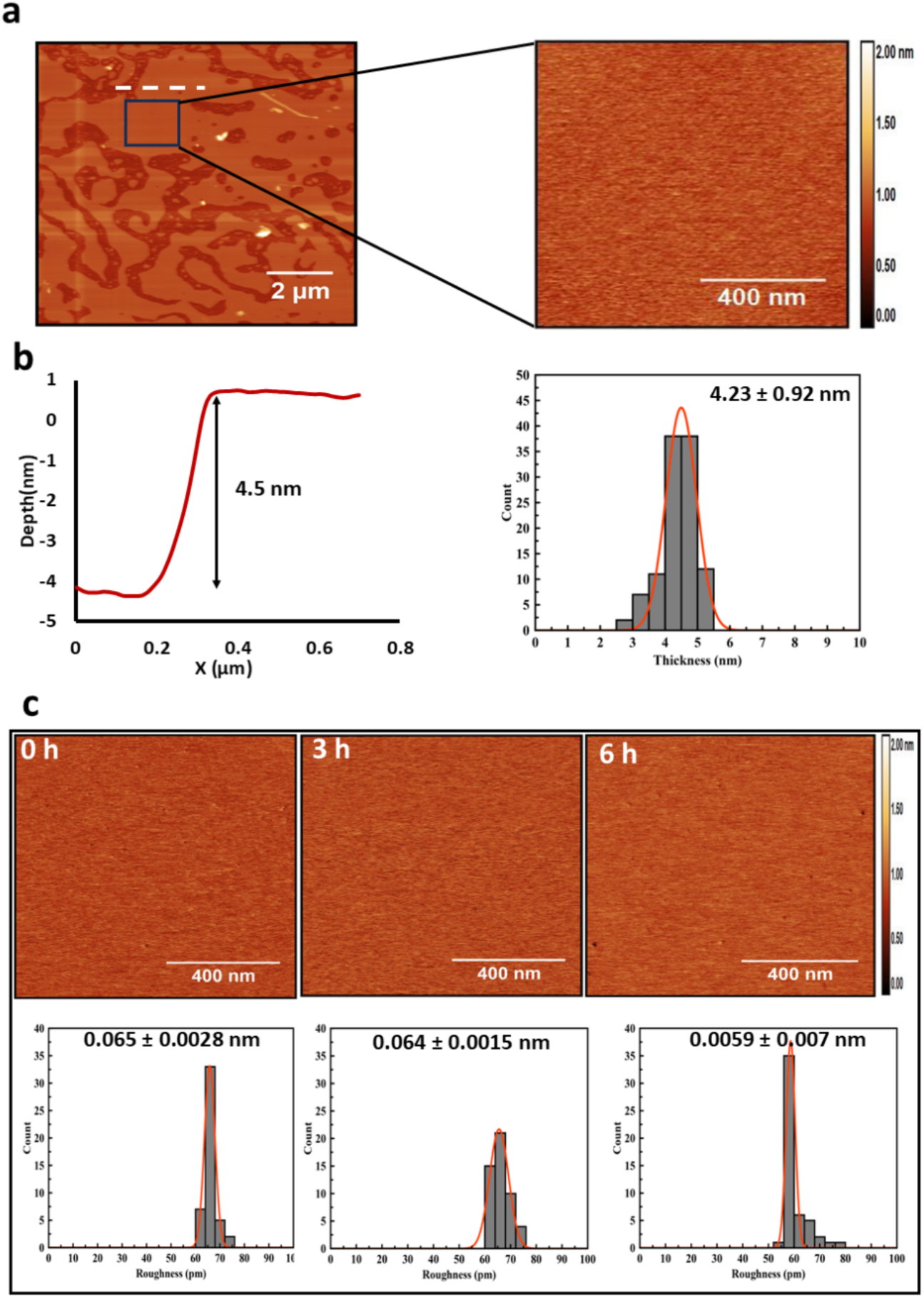
AFM image illustrating the formation of a 0.25 mg/mL POPC:POPS supported lipid bilayer (SLB) incorporating cholesterol, with the corresponding edge height. **a,** AFM image of size 10 x 10 µm scan of POPC:POPS:20% Chol SLB showing appropriate surface coverage as well as edge. Few white features present on the surface are the unruptured vesicles of cholesterol. the zoomed area shows the AFM image of 1 x 1 µm scan of the smooth surface of the SLB. In all the assembled SLB with different mol% of cholesterol, similar surface of 1 x 1 µm was selected for time-lapse AFM imaging until 6 hours. **b,** Height of the edge of the SLB which was measured as 4.5 nm. The cross section was performed on the edge which is shown with white dashed line in fig. a. Several cross-sections was performed on the edge of the bilayer (n=100). The histograms were approximated with Gaussians. The mean height obtained from the approximations along with the standard deviations is shown on right hand side. **c,** Time-lapse AFM images with corresponding RMS roughness of 0.25 mg/mL POPC:POPS:20% Chol bilayer (n=50). The measurement was performed at 0 h, 3 h and 6 h of 1 x 1 µm smooth area of bilayer. The histograms were approximated with Gaussians. The mean RMS roughness obtained from the approximations along with the standard deviations are shown inside each frame.

**Supplementary Fig. 2:**
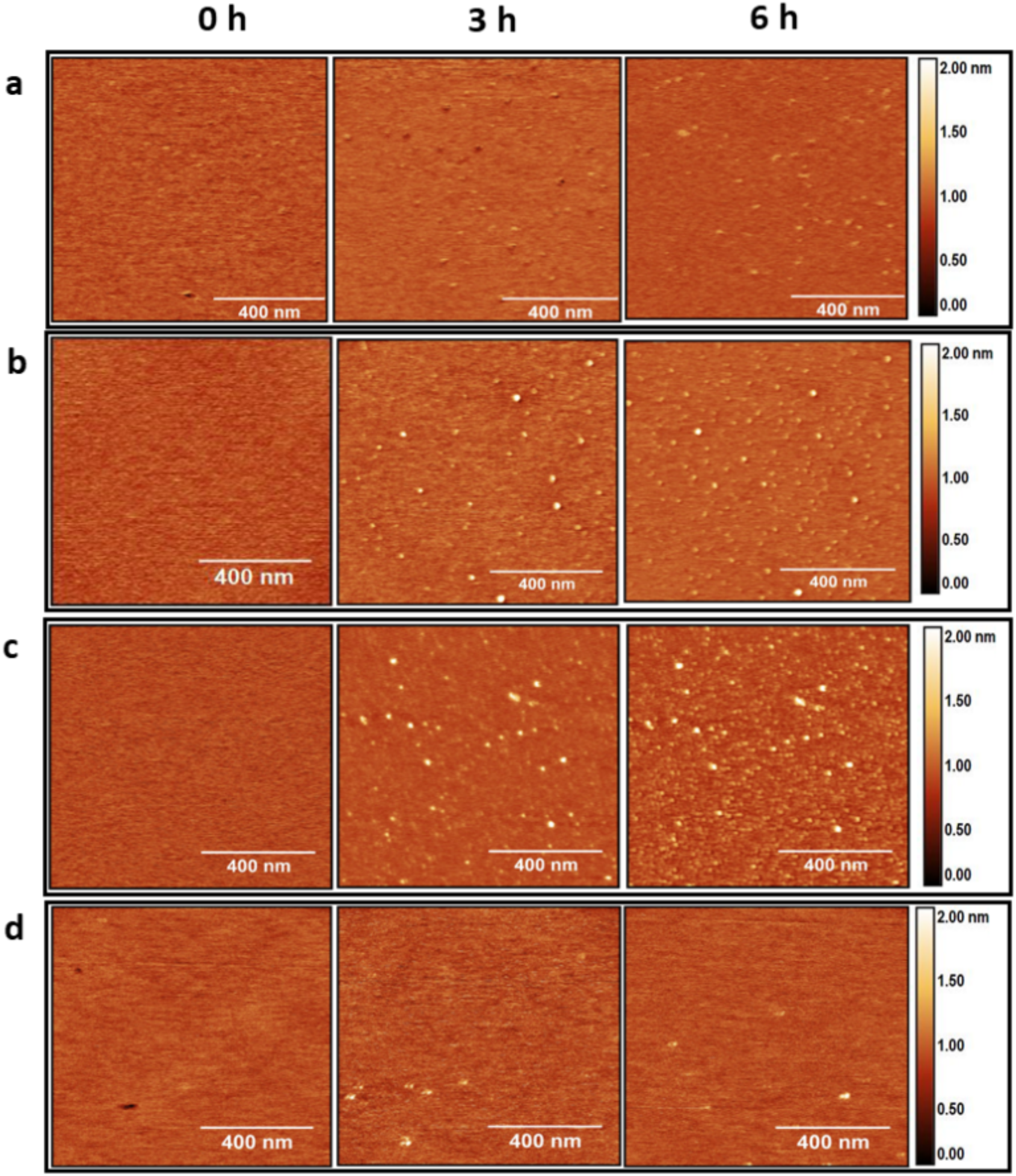
AFM time-lapse images illustrating the aggregation of 50 nM Aβ42 on POPC:POPS bilayer with and without Cholesterol. Aggregation of 50 nM Aβ42 on 0.25mg/mL POPC:POPS lipid bilayer until 6 hour; **a,** with 10% cholesterol **b,** with 20% cholesterol and **c,** with 30% cholesterol **d,** without cholesterol. The scan size is 1 x 1 µm. The number and sizes of the aggregates exhibit a progressive increase over time, with the cholesterol concentration being the highest for the bilayer with 30% cholesterol.

**Supplementary Fig. 3:**
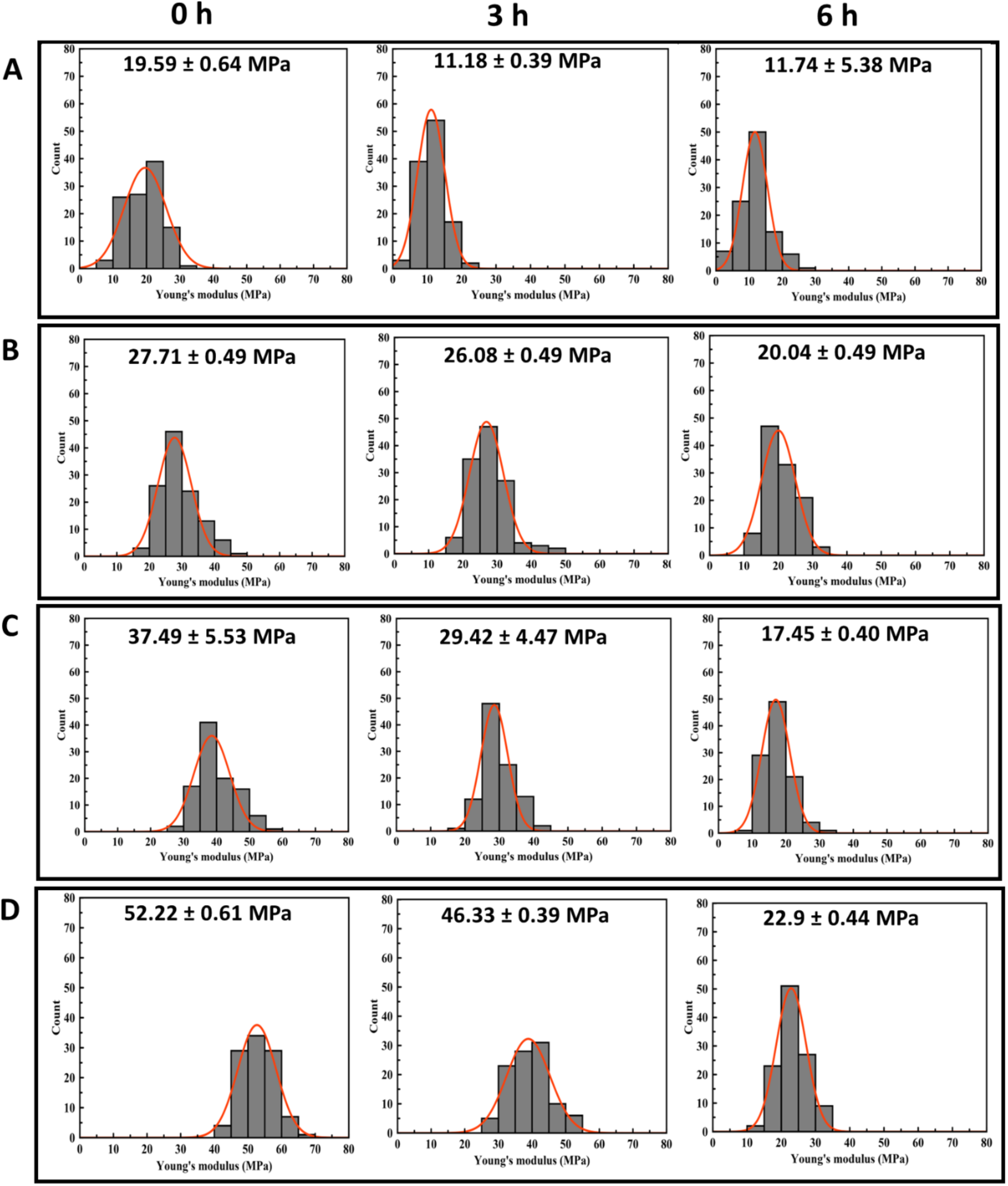
Histogram for values of Young’s modulus of POPC:POPS phospholipid bilayer over 6-hour duration with different mol% of Cholesterol. The Young’s modulus of 0.25 mg/mL POPC:POPS bilayer with **a,** POPC:POPS **b,** POPC:POPS:10% Chol **c,** POPC:POPS:20% Chol **d,** POPC:POPS:30% Chol from 0 time point to 6 h in the presence of 50 nM Aβ42 (n=100) was measured. The modulus data obtained from each force points of the scanned surface were approximated with Gaussians. The mean values obtained from the approximations along with the standard deviations are shown inside each frame.

**Supplementary Fig. 4:**
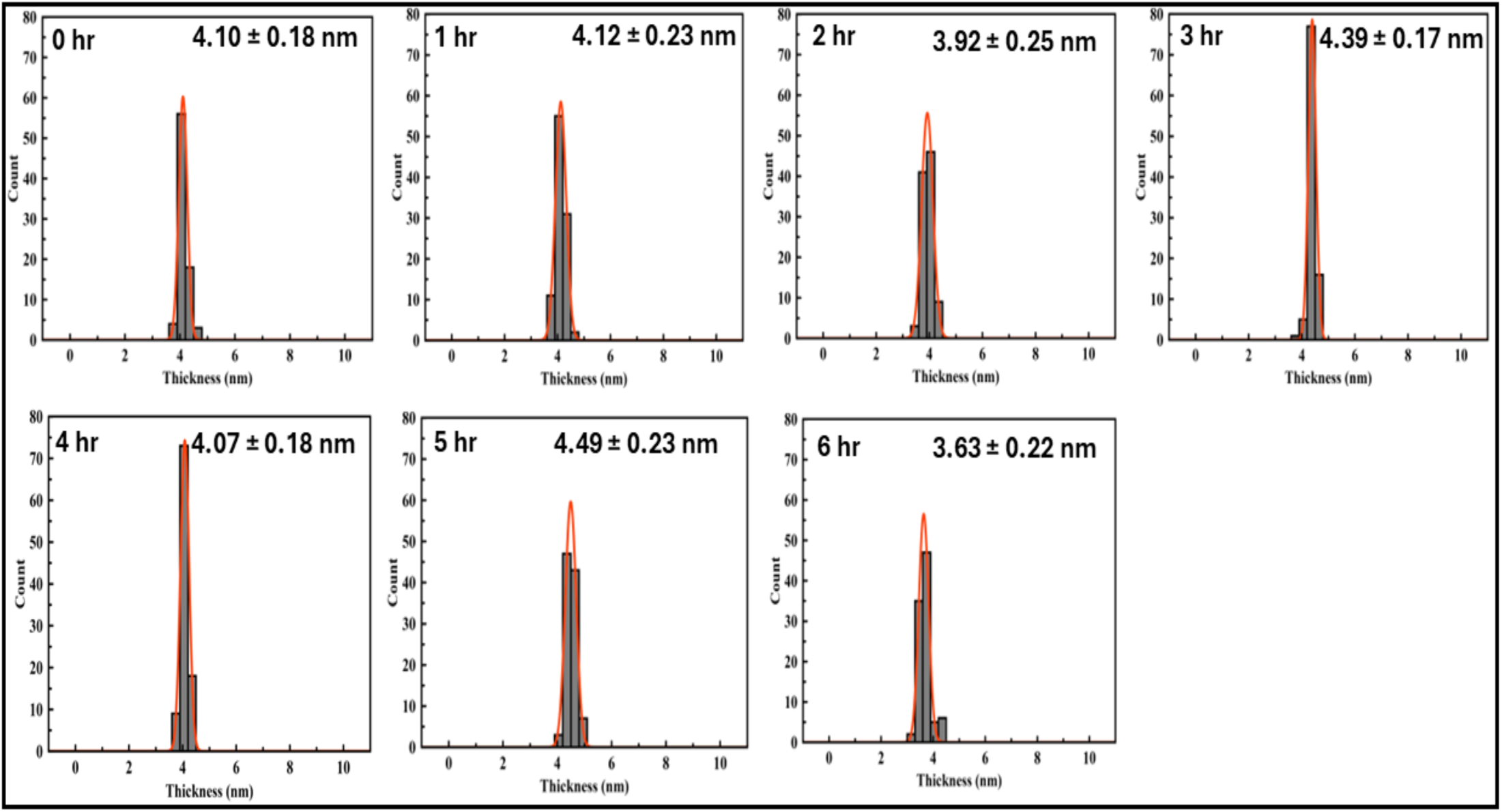
Histograms for thickness of bilayer at each hour over a 6 h duration. The thickness of the bilayer POPC:POPS with 20% Chol with 50 nM Aβ42 above the surface was measured at various locations over the smooth area using the force curves shown in Figure 3B. The histograms were approximated with Gaussians. The mean values obtained from the approximations along with the standard deviations are shown inside each frame.

**Supplementary Fig. 5:**
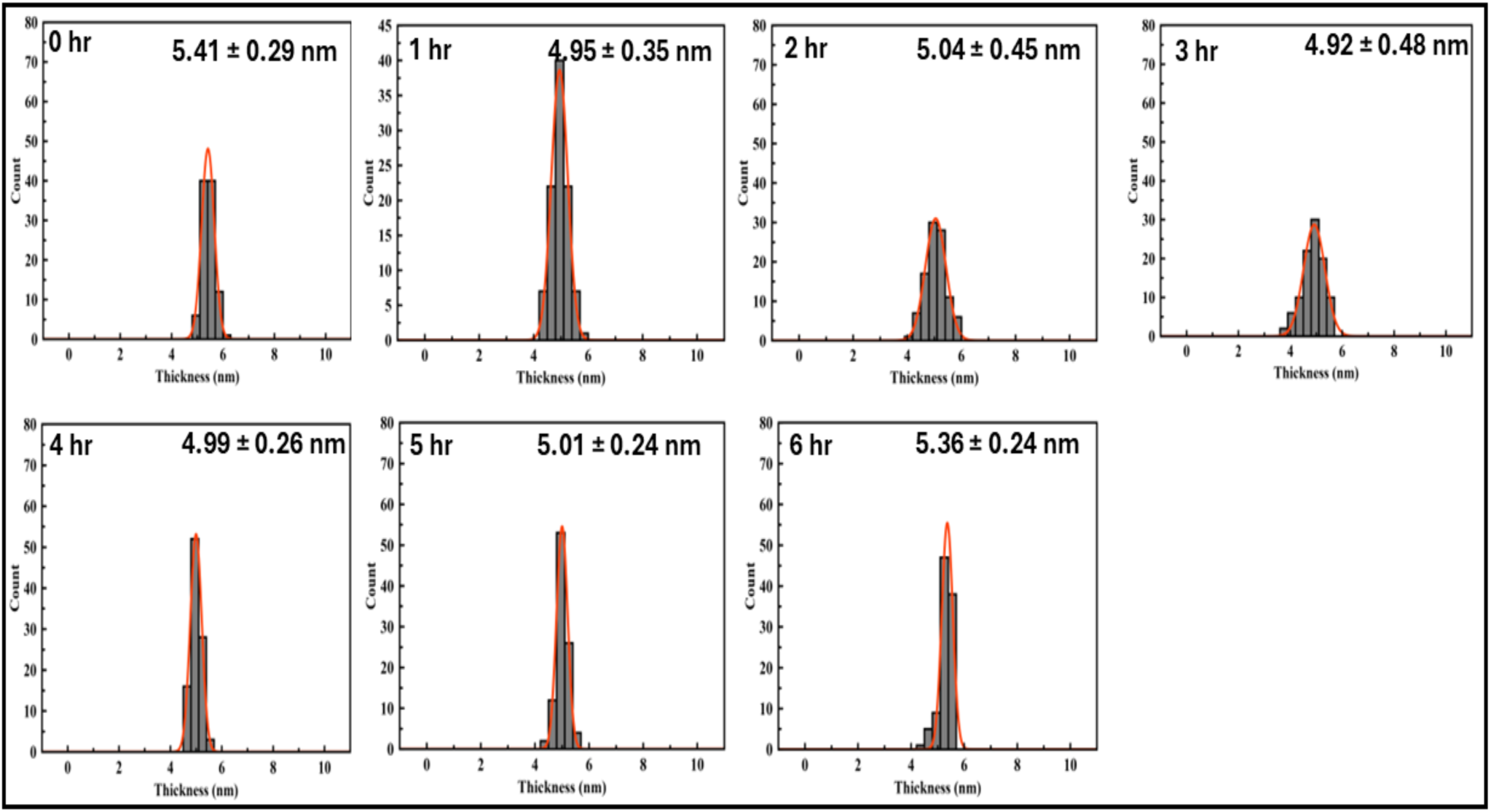
Histograms for values of thickness of bilayer treated with MβCD over a 6 h duration. The thickness of the bilayer POPC:POPS with 20% Chol with 3 Mm MβCD above the surface was measured at various locations over the smooth area. The histograms were approximated with Gaussians. The mean values obtained from the approximations along with the standard deviations are shown inside each frame.

**Supplementary Fig. 6:**
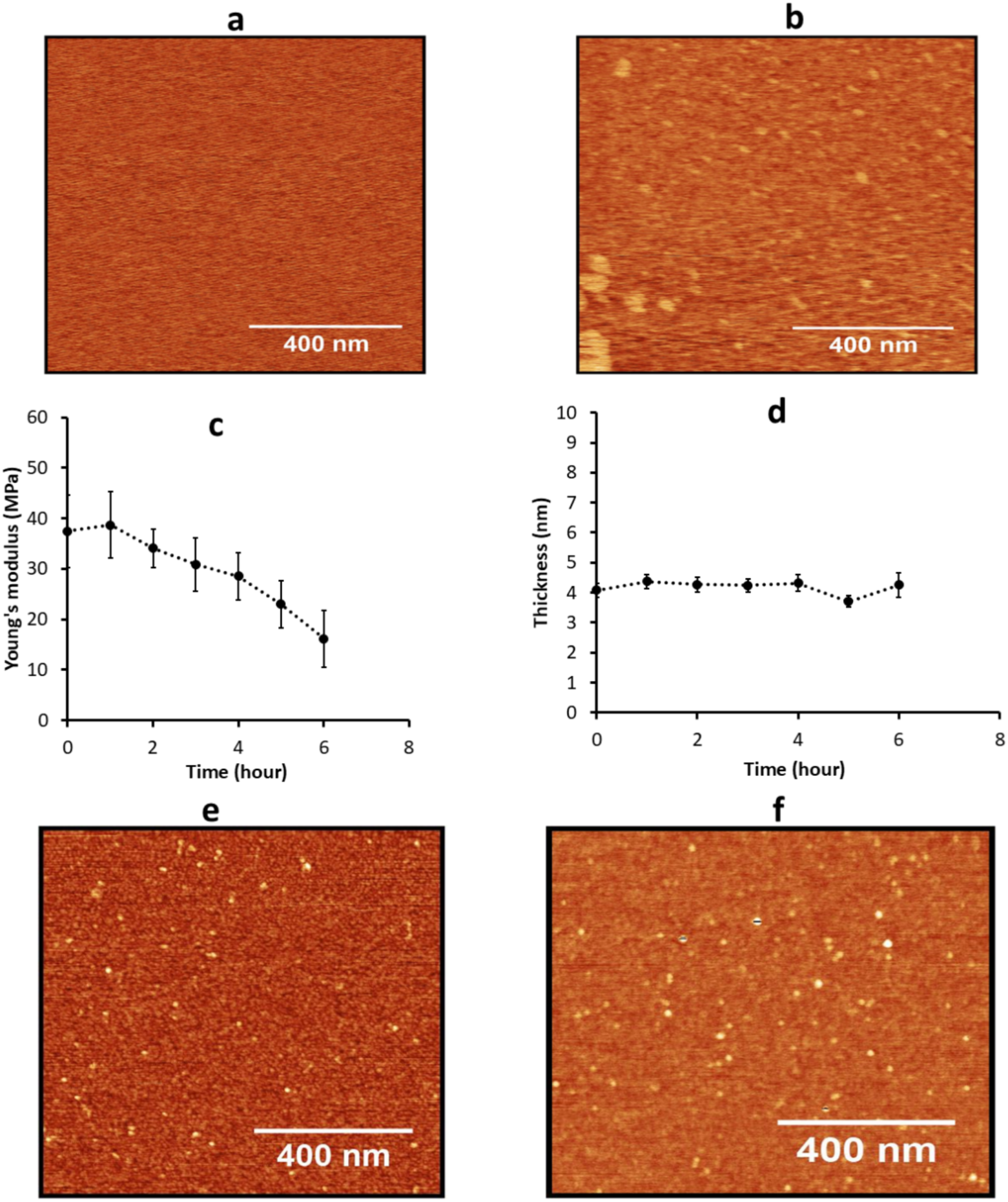
AFM-time lapse imaging and mechanical characterization of bilayer with 25- NBD cholesterol. AFM imaging was performed on a 0.25 mg/mL POPC:POPS phospholipid bilayer incorporating 20% 25-NBD cholesterol in the presence of 50 nM Aβ42, with scans acquired at **a,** 0 hour and **b,** 6 hours. The scan area was set to 1 × 1 µm. **c,** Young’s modulus of the phospholipid bilayer under these conditions was determined (n = 100). **d,** Membrane thickness variations over time (0 to 6 hours) were quantified using force-distance curve analysis. Multiple measurements were conducted across different regions of the 1 × 1 µm scan area, and 100 data points were fitted to Gaussian distributions. The mean thickness values, derived from these approximations, are presented alongside their respective standard deviations as scattered data points. **e,f,** AFM time-lapse imaging was conducted on a dry sample obtained by withdrawing the supernatant from a 0.25 mg/mL POPC:POPS phospholipid bilayer incorporating 20% 25-NBD cholesterol. Images corresponding to time points **e,** 1 hour and **f,** 6 hours post-solution removal were analyzed.

**Supplementary Table 1:** Roughness (pm) of the bilayer without Aβ42.

| **Time (h)** | **PC:PS** | **PC:PS:10% Chol** | **PC:PS:20% Chol** | **PC:PS:30% Chol** |
| --- | --- | --- | --- | --- |
| **0** | 59.47 ± 1.54 | 68.86 ± 4.32 | 67.41 ± 3.56 | 79.16 ± 2.54 |
| **1** | 58.44 ± 0.86 | 67.33 ± 3.11 | 67.65 ± 2.30 | 78.16 ± 1.21 |
| **2** | 59.04 ± 0.42 | 72.44 ± 8.44 | 69.42 ± 3.21 | 81.82 ± 1.01 |
| **3** | 59.33 ± 0.94 | 66.52 ± 4.44 | 73.42 ± 5.72 | 76.52 ± 2.02 |
| **4** | 60.09 ± 0.43 | 68.76 ± 1.95 | 74.7 ± 2.28 | 73.44 ± 1.51 |
| **5** | 60.08 ± 0.54 | 69.91 ± 6.97 | 71.24 ± 6.10 | 96.12 ± 1.17 |
| **6** | 62 ± 1.12 | 70.92 ± 7.42 | 71.36 ± 2.99 | 90.68 ± 3.02 |

Data are expressed as Mean ± SD (n=10)

**Supplementary Table 2:** Young’s Modulus (MPa) of the bilayer with Aβ42.

| **Time (h)** | **PC:PS** | **PC:PS:10% Chol** | **PC:PS:20% Chol** | **PC:PS:30% Chol** |
| --- | --- | --- | --- | --- |
| **0** | 20.12 ± 4.84 | 27.84 ± 5.78 | 38.64 ± 5.21 | 51.14 ± 4.34 |
| **1** | 19.59 ± 6.4 | 27.71 ± 4.9 | 37.49 ± 5.3 | 52.22 ± 6.1 |
| **2** | 12.97 ± 3.2 | 28.42 ± 4.3 | 37.38 ± 5.6 | 52.68 ± 5.7 |
| **3** | 11.18 ± 3.9 | 26.08 ± 4.9 | 29.42 ± 4.7 | 46.33 ± 3.9 |
| **4** | 10.87 ± 3.4 | 20.47 ± 4.3 | 27.04 ± 3.8 | 38.88 ± 6.4 |
| **5** | 9.77 ± 3.6 | 21.36 ± 6.5 | 18.31 ± 3.7 | 28.49 ± 3.6 |
| **6** | 11.74 ± 3.8 | 20.04 ± 4.9 | 17.45 ± 4.0 | 22.9 ± 4.4 |

Data are expressed as Mean ± SD (n=100)

